# A read count-based method to detect multiplets and their cellular origins from snATAC-seq data

**DOI:** 10.1101/2021.01.04.425250

**Authors:** Asa Thibodeau, Alper Eroglu, Nathan Lawlor, Djamel Nehar-Belaid, Romy Kursawe, Radu Marches, George A. Kuchel, Jacques Banchereau, Michael L. Stitzel, A. Ercument Cicek, Duygu Ucar

## Abstract

Similar to other droplet-based single cell assays, single nucleus ATAC-seq (snATAC-seq) data harbor multiplets that confound downstream analyses. Detecting multiplets in snATAC-seq data is particularly challenging due to its sparsity and trinary nature (0 reads: closed chromatin, 1: open in one allele, 2: open in both alleles), yet offers a unique opportunity to infer multiplets when >2 uniquely aligned reads are observed at multiple loci. Here, we implemented the first read count-based multiplet detection method, ATAC-DoubletDetector, that detects multiplets independently of cell-type. Using PBMC and pancreatic islet datasets, ATAC-DoubletDetector captured simulated heterotypic multiplets (different cell-types) with ∼0.60 recall, showing ∼24% improvement over state of the art. ATAC-DoubletDetector detected homotypic multiplets with ∼0.61 recall, representing the first method to detect multiplets originating from the same cell type. Using our novel clustering-based algorithm, multiplets were annotated to their cellular origins with ∼85% accuracy. Application of ATAC-DoubletDetector will improve downstream analysis of snATAC-seq.

## MAIN

Single nucleus ATAC-seq (snATAC-seq)^1–3^ technology is widely used to study epigenomes of diverse cells and tissues with increased resolution^3,4^. However, as with other droplet based single cell technologies, snATAC-seq data harbor multiplet nuclei^5^. The presence of multiplets can confound downstream analyses by introducing combined epigenomic profiles that originate from two or more nuclei, increasing the difficulty of clustering and comparing different cell types within a sample. Compared to other single cell assays, the difficulty of detecting multiplets in snATAC-seq is further increased due to data sparsity and the trinary nature of chromatin accessibility levels (e.g., 0 reads: closed chromatin, 1: open in one allele, 2: open in both alleles).

The current state of the art for detecting multiplets in snATAC-seq data adapt detection methods developed for single cell RNAseq (scRNA-seq). Notably, two snATAC-seq data analysis packages, SnapATAC^6^ and ArchR^7^, either employ or implement a method similar to multiplet detection methods (i.e., DoubletFinder^8^ and Scrublet^9^) for scRNA-seq. In these methods, synthetic heterotypic multiplets (i.e., originating from different cell types) are simulated by combining profiles of two or more cells, which are then used to detect putative multiplets based on cluster similarity. Such algorithms assume that multiplets and singlets exhibit distinct genomic profiles, which becomes problematic when true singlets share genomic profiles with two or more cell types. Under this assumption, these methods will fail to detect homotypic multiplets (i.e., originating from the same cell type) since their overall genomic profile is considered to be similar to that of the underlying cell type. However, homotypic multiplets are characterized by increased read counts compared to singlets, suggesting new methods that utilize read counts can detect them. In order to overcome the limitations of existing methods to detect both homotypic and heterotypic multiplets, we developed a novel multiplet detection method, ATAC-DoubletDetector, that exploits read count distributions to infer multiplets in snATAC-seq data.

ATAC-DoubletDetector’s efficacy was tested in two snATAC-seq datasets generated from peripheral blood mononuclear cells (PBMCs) samples (n=2) and pancreatic islet (n=2) tissues. We identified multiplets in these tissues and quantified the algorithm’s efficacy using simulated homotypic and heterotypic multiplets. We found that when snATAC-seq samples were adequately sequenced (e.g., >20k valid read pairs per cell), ATAC-DoubletDetector proved very effective for detecting both homotypic and heterotypic multiplets (recall ranging from 0.74-0.89 in PBMCs). In addition, ATAC-DoubletDetector includes a novel clustering-based algorithm that accurately annotates the cellular origins of detected multiplets (85% average accuracy in our simulations), providing further data quality insights. ATAC-DoubletDetector is provided as a user-friendly computational framework with documentation and source code freely available at: https://github.com/UcarLab/ATAC-DoubletDetector.

## Results

ATAC-DoubletDetector leverages the fact that the expected number of uniquely aligned reads for a given locus ranges from 0 to 2 per nucleus in snATAC-seq data: 0 = closed chromatin, 1 = open in one allele (i.e., from either maternal or paternal chromosomes), 2 = open in two alleles (i.e., both maternal and paternal chromosomes) (Fig. 1a). A locus can have more than two reads (>2) when: 1) it contains repetitive sequences; 2) there are sequencing or alignment errors; or 3) reads stem from multiplet nuclei. In the case of multiplets, we expect to observe many loci with >2 reads since their epigenomic profiles are derived from two or more nuclei resulting in increased accessible DNA. ATAC-DoubletDetector identifies all loci with >2 reads for each cell/nucleus (Fig. 1b) by utilizing sorted read alignments to detect their overlapping read intervals (22-39 bp on average across all samples). A unified list of these loci across all nuclei is then generated to quantify the number of occurrences where >2 reads align to a locus in a given nucleus (Fig. 1c). As a proof of concept, highly significant multiplets (P-Values < 10^−324^) can be clearly seen harboring many more loci with >2 reads (924-1054 loci) than average (∼23 loci per nuclei) (Extended Data Fig.1). Random occurrences of loci with >2 reads (i.e., due to sequencing or alignment errors) were modeled with the Poisson cumulative distribution function using the mean number of overlaps detected across all cells. Nuclei that harbor significantly more loci with >2 reads are identified as multiplets based on their deviations from the distribution using False Discovery Rate (FDR) (Fig. 1c). To trace multiplets back to their cellular origins, we employed a clustering-based algorithm as part of the ATAC-DoubletDetector framework. Marker peaks are detected to generate reference accessibility profiles for each cell type using single cell clustering. Epigenomic similarity scores at marker peaks are then used to compare multiplet profiles with singlet profiles to differentiate between heterotypic and homotypic multiplets and annotate them.

**Fig 1:**
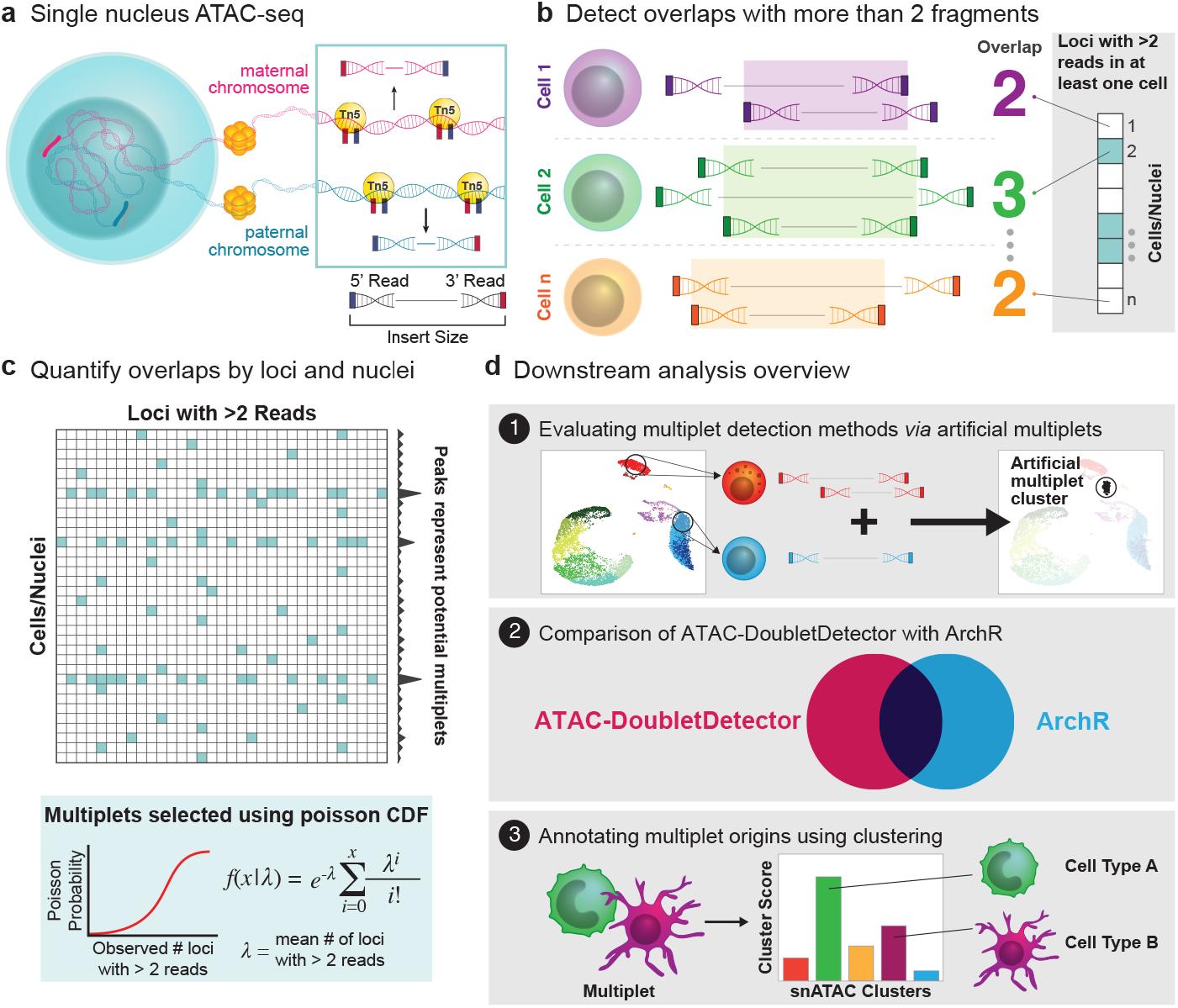
Overview of detecting multiplets in snATAC-seq. **a**, Tn5 transposase cleaves accessible DNA at maternal and paternal chromosomes. Number of ATAC-seq read counts per loci per nucleus are expected to be 0, 1, or 2. **b**, Instances where more than 2 (>2) reads are observed for any locus in a cell are identified using an efficient algorithm for counting the number of overlapping reads. **c**, Poisson cumulative distribution function is used to detect multiplets based on deviations from expected number of loci with >2 reads. **d**, Overview of downstream analyses: 1) quantification of multiplet detection performances using artificial multiplets, 2) comparison of ATAC-DoubletDetector to alternative method ArchR, 3) annotating cellular origins of multiplets using a clustering-based method.

We demonstrate the utility and performance of our computational framework by applying our methods in PBMC and islet sample datasets (Fig. 1d). First, we simulated artificial multiplets in PBMC and islet samples and quantified ATAC-DoubletDetector’s ability to identify and annotate these multiplets. Second, we compared ATAC-DoubletDetector to ArchR, measuring their overall performances and their ability to detect simulated heterotypic and homotypic multiplets. Finally, we measure the efficacy of our annotation method and analyze multiplet cellular origins to understand whether cell type influences the rate of multiplet occurrences.

### ATAC-DoubletDector detects heterotypic and homotypic multiplets in PBMC and islet samples

We generated snATAC-seq libraries from two human PBMC and two human pancreatic islet samples using 10x Genomics Chromium platform^3^. Sequence reads were preprocessed using Cell Ranger ATAC pipeline (methods), resulting in an average of 5,559 and 6,173 nuclei per sample and an average of 24,393 and 16,625 valid read pairs per cell for PBMC and islet samples respectively (Fig. 2a). Valid read pairs refer to all pairs of paired end reads that align to autosomes and pass quality control flags/thresholds (methods). Despite deeper sequencing for islet samples, fewer valid read pairs were observed in islet samples compared to PBMC samples (Fig. 2b), which can be explained by increased mitochondrial reads in islets (114,821,502 and 47,522,248 total reads aligned to chrM) compared to PBMCs (2,610,761and 947,233 total reads aligned to chrM).

**Fig 2:**
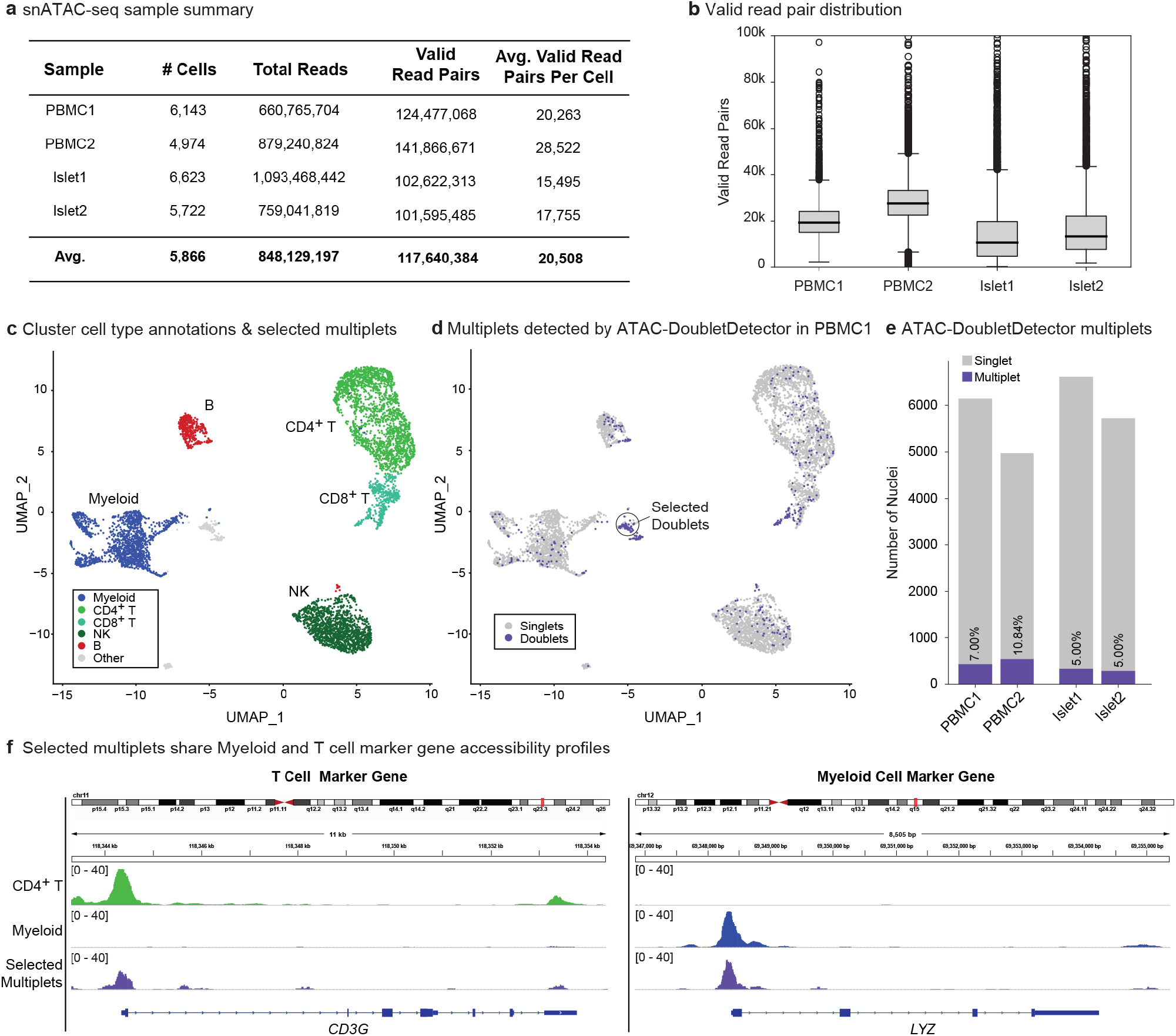
ATAC-DoubletDetector identifies heterotypic and homotypic multiplets in human PBMC snATAC-seq data. **a**, Summary of snATAC-seq samples generated and used in this study from human PBMC and islets. **b**, Valid read pair distributions for PBMC and islet snATAC-seq samples. **c**, PBMC clusters were annotated based on their correlations with sorted bulk ATAC-seq data (See. Extended Data Fig.2). **d**, All multiplets (heterotypic and homotypic) detected by ATAC-DoubletDetector in PBMC1. Selected multiplets refer to multiplets for which aggregated profiles are shown in panel **f** of this figure. **e**, The number of cells and percentage of multiplets detected by ATAC-DoubletDetector in PBMC and islet samples. **f**, Chromatin accessibility profiles of CD4^+^ T, myeloid, and selected multiplets around for T cell marker gene (*CD3G*) and myeloid cell marker gene (*LYZ*). CD4^+^ T and myeloid cells show strong accessibility signals for their relevant marker genes while selected multiplets have accessible chromatin for both marker genes.

Nuclei clustering using an in-house implementation (methods) of a two-pass clustering method^3^ for snATAC-seq data identified 16 and 15 clusters for PBMC1 and PBMC2. Correlating pseudo-bulk accessibility profiles of these clusters with accessibility maps from sorted bulk ATAC-seq data^10^ (Extended Data Fig. 2a,b) grouped them into 5 major cell types: myeloid (including CD14+, CD16 monocytes and conventional dendritic cells), B, CD4^+^ T, CD8^+^ T, and NK cells (Extended Data Fig. 2c,d). These annotations were confirmed based on chromatin accessibility patterns at cell-specific marker genes (Extended Data Fig. 3a,b). The same clustering procedure identified 14 and 12 distinct clusters for islet1 and islet2, which were then annotated as alpha, beta, delta, and ductal cells by integrating their accessibility profiles with in-house islet scRNA-seq data (Extended Data Fig. 4a,b). These annotations were confirmed by analyzing the chromatin accessibility patterns at known cell-specific marker genes^11^ (Extended Data Fig. 4c,d).

We applied ATAC-DoubletDetector on PBMCs and human islet samples using an FDR cutoff of 0.01 (Methods). Nuclei detected as multiplets were distributed throughout all clusters (Fig. 2c-d, Extended Data Fig. 5) and in one case (PBMC1) multiplets formed their own distinct cluster (see selected multiplets in Fig. 2d). The percentage of detected multiplets were higher in PBMCs (7%, 10.84%) compared to islets (5% for both samples) (Fig. 2e), which is likely due to the lower valid read pairs per nuclei in islets as previously mentioned (Fig. 2b).

To further study the biological relevance of these detected multiplets, we selected a cluster which exclusively encompassed multiplets (Fig. 2d; PBMC 1 selected multiplets) and analyzed their chromatin accessibility profiles (Fig. 2f). The selected multiplets were characterized by a high chromatin accessibility at the promoters of both *CD3G* (T cell marker gene) and *LYZ* (monocyte marker gene), suggesting T cell-monocyte multiplets. These results demonstrate how read count distribution information from snATAC-seq can be used to effectively detect multiplets.

### ATAC-DoubletDetector effectively detects simulated heterotypic and homotypic multiplets

To quantify the efficacy of ATAC-DoubletDetector, we generated artificial multiplets by randomly selecting 5% of nuclei in a sample and pairing them together to artificially form multiplets (repeated 10 times per sample). This resulted in artificial multiplets at 2.5% of the total number of nuclei within a sample. These artificial multiplets serve as positive multiplet examples and enable us to measure recall (i.e., the fraction of detected artificial multiplets among all artificial multiplets introduced in the sample). We first evaluated ATAC-DoubletDetector’s ability to detect heterotypic, homotypic, and a combination of both multiplet types. We then compared it’s performance in comparison to another method ArchR^7^.

ATAC-DoubletDetector detected heterotypic multiplets introduced in PBMC samples with high recall (average recall 0.80 for PBMC1 and 0.90 for PBMC2 over 10 runs), outperforming ArchR (0.23 and 0.24 respectively) (Fig. 3a). Average recall for ATAC-DoubletDetector was lower in islet1 and islet2 than PBMCs (0.37 and 0.34 average recall respectively) whereas the average recall showed improvement for ArchR (0.68 and 0.30 average recall respectively). Decreased performance of ATAC-DoubletDetector’s in islets can be explained by low number of valid read pairs per nuclei in islet samples compared to PBMCs (Fig 2b). Notably, ATAC-Doublet detector was equally effective for detecting homotypic multiplets (average recall 0.82 and 0.91 for PBMC 1 and PBMC 2, 0.38 and 0.31 for islet 1 and islet 2) (Fig. 3b), demonstrating the utility of using read counts to detect multiplets. As expected, ArchR had low recall for detecting homotypic multiplets (average between 0.07 and 0.11 for all samples), as this algorithm identifies multiplets with distinct genomic profiles from singlets. Finally, we measured the efficacy to simultaneously detect both types of multiplets by introducing a more realistic-heterotypic and homotypic multiplet 1:1 ratio (Extended Data Fig. 6a). As expected, the average recall values of ATAC-DoubletDetector’s were similar (0.82 and 0.92 for PBMC1 and PBMC 2, 0.34 and 0.33 for islet1 and islet2 respectively), while, those of ArchR were lower (0.13 and 0.16 for PBMC1 and PBMC2, 0.40 and 0.17 for islet1 and islet2), likely due to its poor homotypic multiplet detection performance.

**Fig 3:**
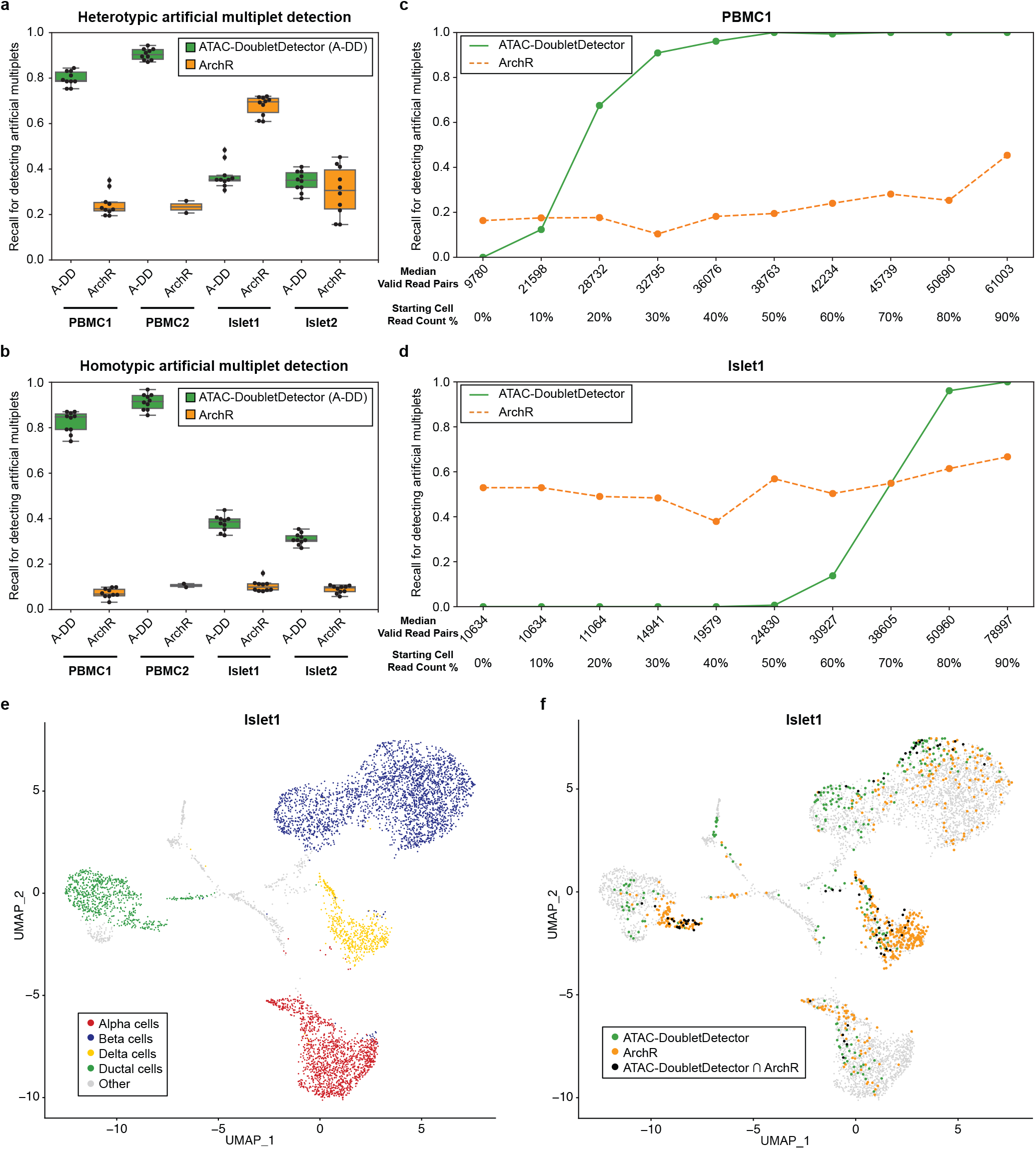
ATAC-DoubletDetector detects multiplets with high recall when read depth is sufficient. **a**-**b**, Recall for detecting heterotypic (**a**) and homotypic (**b**) artificial multiplets. ATAC-DoubletDetector consistently detected both heterotypic and homotypic multiplets with similar recall, while ArchR was only effective for predicting heterotypic multiplets for data with high heterogeneity. **c-d**, Performance of detecting artificial multiplets at increasing valid read pair (insertions) distributions for PBMC1(**c**) and islet1(**d**). ATAC-DoubletDetector effectively detects multiplets at the >40k valid read pairs per nucleus. ArchR’s performance did not observe the same level of effect for read depth. **e**, Reference annotations for islet1. Islet1 annotations correspond to alpha, beta, delta and ductal cell types. **f**, Representative UMAP plots for multiplets detected by ATAC-DoubletDetector and ArchR for islet1 (other samples shown in Extended Fig. 8). We identified islet clusters for Alpha, Beta, Delta, and Ductal cells. Majority of multiplets detected were not shared between the two methods. Heterotypic multiplets were the most common. Note: ArchR detected the majority of Delta cells as multiplets.

To further study how the valid read pairs influence ATAC-DoubletDetector’s performance, we generated artificial multiplets using cells with ranging reads per nucleus (Fig 3c-d, Extended Data Fig. 6b). We observed a noticeable increase in average recall (> 0.96 recall) for ATAC-DoubletDetector, when the number of valid read pairs was above 47.2k, corresponding to an average of 23.6k valid reads pairs per nucleus. In contrast, ArchR did not show significant differences in performances with respect to the number of valid read pairs per nucleus (Extended Data Fig. 6b), as it relies more on genomic profile similarity to detect multiplets. More exhaustive analyses of 100 repetitions per sample further confirmed that the majority (96%, 98% for PBMC1 and PBMC2 and 83%, 72% for islet1 and islet2) of multiplets with >40k valid read pairs (i.e., multiplets formed from nuclei with 20k valid read pairs each) were detected with this method (Extended Data Fig. 7). Together, these analyses suggest that when >20k valid read pairs are captured per nucleus, ATAC-DoubletDetector is very effective in detecting both homotypic and heterotypic multiplets from snATAC-seq data.

To compare ATAC-DoubletDetector and ArchR performances, we ran ArchR with recommended parameter settings (i.e., k=10 nearest neighbors and 1.5 filter ratio). Only 38 to 78 multiplets across all samples were detected by both methods (Fig 3e-f, Extended Data Fig. 8, Extended Data Fig. 9a-b) and majority of these multiplets were among the ones that formed their own clusters (i.e., heterotypic multiplets). For example, the majority of selected multiplets detected in cluster in Fig 2d were detected by both methods (Extended Data Fig. 8), which are multiplets that have unique epigenomic profiles; hence easier to detect with the synthetic multiplet-based method employed by ArchR. Notably, 47.35% of Delta cells were identified as multiplets by ArchR for Islet1 (Figure 3f, Extended Data Fig. 8). Delta cells resemble both alpha and beta cells in their genomic profile, hence these cells were mistakenly detected as multiplets by ArchR, demonstrating a pitfall for synthetic multiplet-based methods. Multiplets are expected to have higher read counts than singlets since they combine chromatin accessibility profiles of more than one nucleus. In alignment with this, multiplets detected by ATAC-DoubletDetector had significantly higher valid read pair counts compared to singlets (average valid read pairs of 46,980 for multiplets and 18,561 for singlets for all samples) (P-Values < 1.375 × 10^−152^). In contrast, read counts for ArchR multiplets were significantly lower (average P-Values < 1.016 × 10^−57^) than ATAC-DoubletDetector multiplets, observing read counts closer to that of singlets (average read count per cell 23,703 for ArchR multiplets and 19,951 for singlets) (Extended Data Fig. 9c). In summary, these analyses showed that when there is sufficient number of valid read pairs per cell (> 20k), count based methods are advantageous over synthetic multiplet-based methods as they can accurately detect both homotypic and heterotypic multiplets.

### Marker peaks can effectively annotate cellular origins of multiplets

Cellular origin annotations of multiplets were inferred using a three-step algorithm (Fig. 4a). First, nuclei were clustered and annotated to their respective cell types. Second, marker peaks were detected for each cluster/cell type. Third, we calculated epigenomic similarity of each multiplet to different cell types by counting marker peak reads for the multiplet and the k=15 nearest neighbor nuclei (Methods). Cluster similarity scores were then used to annotate multiplets. For example, in PBMCs, for each multiplet we calculated 5 scores, where each score represents the similarity of the multiplet epigenome to that of the five studied clusters (Figure 4b). The distribution of these similarity scores are used to first distinguish heterotypic and homotypic multiplets, by comparing their profiles to annotated singlets (Methods). For example, in PBMC1, nuclei in B cell cluster (cluster 5) had high similarity score for B cell marker peaks and low scores for all other cell types (Figure 4b). In contrast, nuclei in cluster 13 had high similarity scores for NK, CD4^+^ T, CD8^+^ T and myeloid cells, a signature of heterotypic multiplets (Fig. 4b). Once the multiplet type is identified, their cellular origins are annotated using the highest scoring cell type(s).

**Fig 4:**
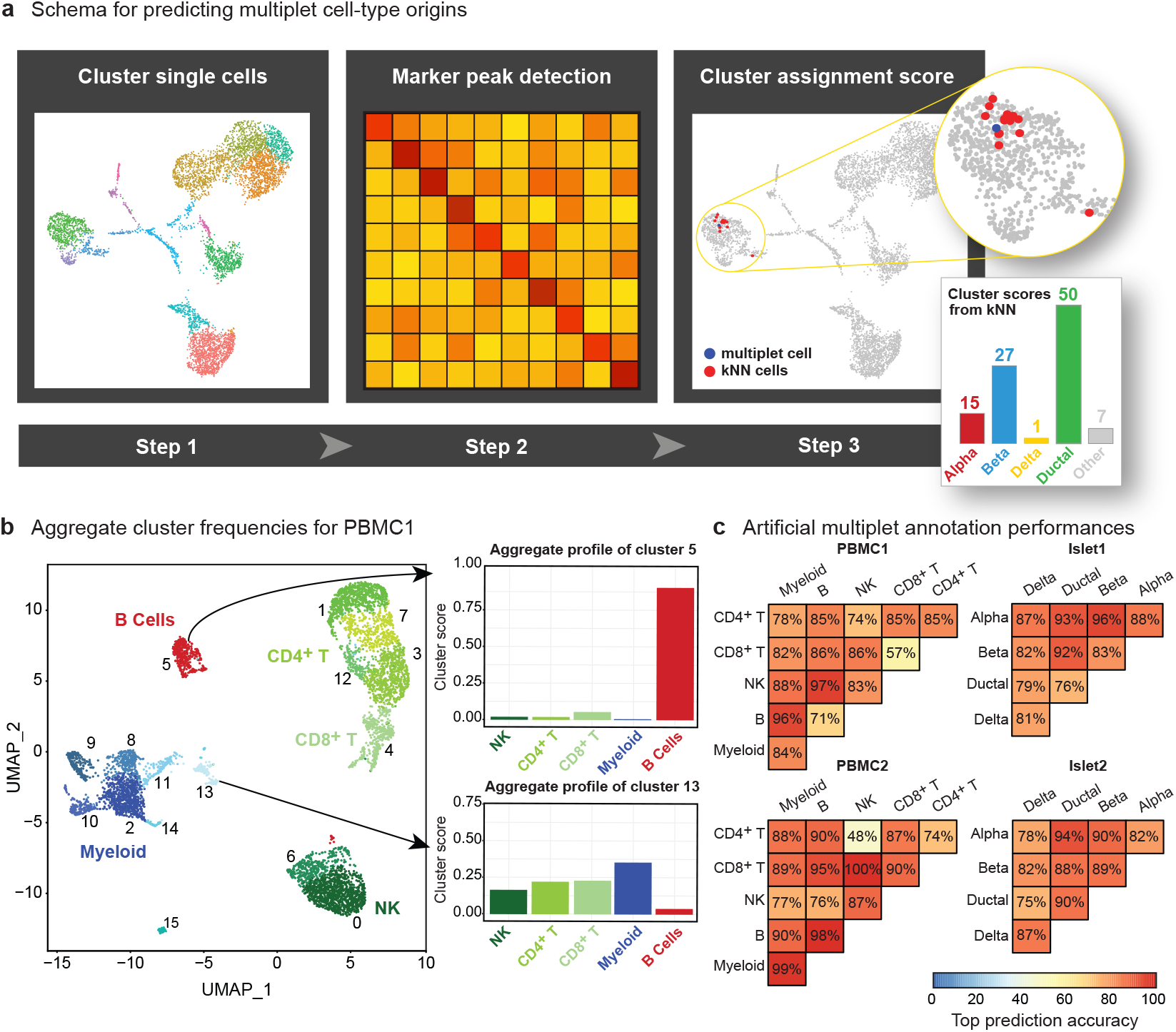
Multiplet cell-type origins are predicted with high accuracy. **a**, Overview of the cell origin annotation pipeline. First, cells are clustered. Second, marker peaks are identified. Third, multiplets and their k-nearest neighbor cells are used to generate cluster similarity scores. **b**, Example of aggregate cluster profiles for predicting cell origin annotations. Clusters corresponding to cell types observe strong signal for their respective cell types (e.g., Cluster 5) while clusters corresponding to multiplets show a mixed profile of cell types (e.g., Cluster 13). **c**, Heatmaps of cell origin annotation accuracies for predicting artificial multiplets derived from cells of the specific cell type pairings. Multiplet annotations showed high accuracies for the majority of cell type compositions.

We evaluated the efficacy of this annotation pipeline using artificial multiplets, where cells were randomly selected and paired together to form both heterotypic and homotypic multiplets. Using these artificial multiplets, we categorized multiplets as homotypic or heterotypic and annotated multiplets with respect to the number of cell types associated with them. We identified the cellular origins of both types of multiplets with an average accuracy of 82.47%, 85.87% in PBMC1, PBMC2 and 85.7%, 85.5% in islet1, islet2 (Fig. 4c). For example, in PBMC1, 96% of all simulated B and myeloid multiplets were correctly annotated. Cell types that have similar functions, hence similar epigenomes, observed lower annotation accuracies; such as 86% for simulated NK and CD8+ T cells. Our annotations were equally effective for annotating both homotypic and heterotypic multiplets, showing 83.65% accuracy on average to annotate homotypic multiplets and 85.59% accuracy to annotate heterotypic multiplets.

### Multiplet cell-type compositions reflect cellular compositions of the underlying tissue

Using ATAC-DoubletDetector’s annotation pipeline, we annotated all detected multiplets in PBMCs and islets. Inspection of aggregate accessibility profiles at marker gene promoters (*MS4A1, CD3G, CD4, CD8A, TREM1, NKG7*, and *KLRF1*) for each cell type in PBMC2 (Fig. 5a) revealed that annotated multiplets have accessibility at relevant marker gene promoters. For instance, homotypic B cell multiplets had strong signal at the promoter of B cell marker gene *MS4A1*, whereas heterotypic multiplets originating from CD8^+^ T cell and B cells had high accessibility signals for both B cell marker gene *MS4A1* and CD8^+^ T cell marker gene *CD8A*.

**Fig 5:**
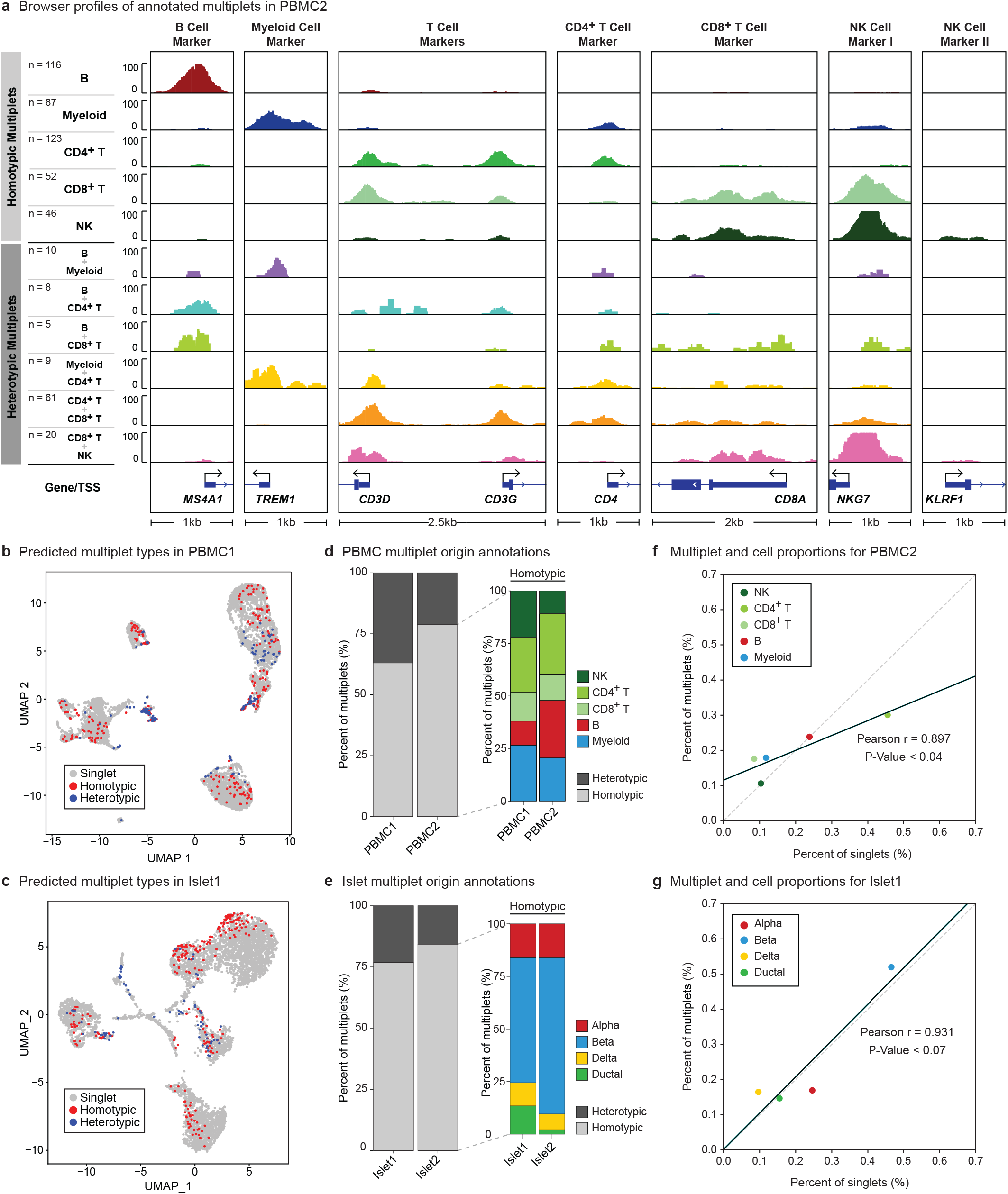
Majority of multiplets are homotypic and correspond to cell type proportions. **a**, Accessibility maps for cell origin annotations for multiplets identified in PBMC2. Homotypic multiplets observe strong signal for their respective marker genes. Heterotypic multiplets observe a combined signal at respective marker genes corresponding to the respective annotated cell types. **b-c**, UMAP clustering for heterotypic and homotypic multiplet annotations in PBMC1 (**b**) and islet1 (**c**). Heterotypic multiplets are found between major cell type clusters. Homotypic multiplets are observed on the periphery of major cell type clusters. **d**-**e**, Heterotypic and homotypic multiplet cell distributions (left bars). Homotypic cell type annotations (right bars) for PBMC (**d**) and islet (**e**) samples. Majority of multiplets are annotated as homotypic. Homotypic cell type distributions show similar distribution to the overall proportions of each cell type in their respective samples. **f**-**g**, Cell and multiplet proportions for PBMC2(**f**) and islet1(**g**). Multiplet cell type proportions are highly correlated with overall cell proportions.

As expected, homotypic multiplets clustered together with the underlying cell type, whereas heterotypic multiplets typically formed their own clusters (Fig. 5b-c, Extended Data Fig. 10a-b). The majority of heterotypic multiplets for islet1 were found between major cell type clusters and near the delta cell cluster while homotypic multiplets resided within the boundaries of singular cell type clusters (Fig. 5d). For PBMC1, the majority of multiplets resided within multiplet cluster we previously identified and as a subcluster of CD8^+^ T cells (Fig. 5e). As before, homotypic multiplets were found within corresponding cell type clusters. Overall, the majority of detected multiplets were homotypic (76.7-84.3% in islets, 63-78.7% in PBMCs), with cell types being distributed with respect to their cell proportions for both homotypic and heterotypic multiplet types (Fig. 5d-e, Extended Data Fig. 10c-d). Indeed, in both tissues, the propensity of a cell type to form a multiplet was positively correlated with the percent of that cell type within the tissue (Pearson’s R = 0.824, 0.897, P-Value < 0.087, 0.04 for PBMC1 and PBMC2, Pearson’s R = 0.931, 0.475 P-Value < 0.07, 0.525 for islet1 and islet2) (Fig. 5f-g, Extended Data Fig. 10e-f), suggesting that snATAC-seq multiplets are more likely to occur randomly than through specific interactions between nuclei. For example, the most abundant cell type in islet1 was beta cells (46.62% of the cell population) which contributed to 51.96% of multiplets (Fig. 5f). Heterotypic multiplet annotations in islet samples mostly originated from alpha, beta and delta cells. In PBMCs, the most frequent heterotypic multiplets were the ones stemming from CD4^+^ T and CD8^+^ T cells (Fig. 5f, Extended Data Fig. 10e).

## DISCUSSION

Detecting and discarding multiplets from snATAC-seq data is a critical step for improving data quality as multiplets can form their own clusters and can confound downstream analyses. ATAC-DoubletDetector exploits read count distributions for a given nucleus to effectively detect and eliminate multiplets without requiring prior knowledge of cell-type information. It accomplishes this by first efficiently counting loci with >2 uniquely aligned reads per nucleus and identifying nuclei with read count distributions deviating from expectations. Unlike other methods that utilize artificial multiplet examples to identify putative multiplets (i.e., ArchR), ATAC-DoubletDetector is capable of detecting both homotypic (i.e., multiplets originating from the same cell type) and heterotypic multiplets (i.e., multiplets originating from different cell types). Eliminating heterotypic multiplets is essential for improved clustering and differential analyses between clusters and samples, whereas homotypic multiplets introduce bias in allele-specific analyses. Hence, detecting and removing both types of multiplets will improve downstream analyses.

The number of valid read pairs per cells is the most important factor affecting the performance of ATAC-DoubletDetector. When read depth per nucleus is sufficiently high (e.g., >20k read pairs per nucleus), ATAC-DoubletDetector is very effective in detecting both heterotypic and homotypic multiplets (average recall = 0.836 to detect artificial multiplets in PBMCs). Since ATAC-DoubletDetector does not depend on artificial multiplet examples, it is not inherently biased towards cell types that resemble others. For example, in islets, delta cells transcriptionally resemble alpha and beta cells, hence artificial multiplets generated by combining alpha and beta cells have genomic profiles that resemble delta cells. These instances are particularly challenging for methods that depend on artificial multiplet examples (e.g., ArchR for snATAC^7^, DoubletFinder^8^ and Scrublet^9^ for scRNA-seq). In alignment with this, ArchR categorized 47.35% of delta cells as multiplets in islet1. Given the success of ATAC-DoubletDetector for identifying multiplets from snATAC-seq data with enough reads per nuclei, it can also be effective in detecting and eliminating multiplets in recent multi-ome transcriptome and epigenome assays^12^.

Epigenomic signal at marker peaks is an effective way to annotate cellular origins of multiplets, where we achieved 84.69% accuracy on average in simulations. Annotations of detected multiplets showed that majority are homotypic. Furthermore, the propensity of nuclei to form multiplets was positively correlated with the abundance of that cell type within the tissue. Since cells are lysed and nuclei are profiled in snATAC-seq protocols^3^; these assays will likely not be prone to biological multiplets due to cell-cell interactions). Therefore, snATAC-seq multiplets likely occur randomly among all cells; hence the most abundant cells are the most likely to form multiplets.

Quantifying the efficacy of multiplet detection methods is a challenging task since true examples of singlet and multiplets are not known. To overcome this challenge, we evaluated ATAC-DoubletDetector’s ability to capture multiplets by simulating artificial multiplets, enabling us to measure recall. ATAC-DoubletDetector identified 5-10.84% of cells as multiplets in islet and PBMC samples, which was in alignment with expectations. Hence, we believe false positive calls are also restricted in our method. Although we quantified our method by forming artificial multiplets, ATAC-DoubletDetector pipeline can be easily extended to capture and annotate multiplets that include data from multiple nuclei.

Multiplets are inevitable in single cell sequencing and performing better data analyses calls for their removal. ATAC-DoubletDetector introduces a novel and effective count-based solution for detecting multiplets and provides a framework for annotating their cellular origins, improving future downstream analyses. ATAC-DoubletDetector code and documentation is freely available at https://github.com/UcarLab/ATAC-DoubletDetector, providing an easy to use interface for all backgrounds. Our multiplet detection algorithm is fast and can be incorporated into data analyses pipelines, where processing of an average library (i.e., ∼5,886 cells at ∼20,508 valid read pairs per cell) takes <30 minutes.

## METHODS

### snATAC-seq cell labeling, capture, library preparation, and sequencing

For single nucleus ATAC sequencing (snATACseq) experiments, viable single cell suspensions from each sample were used to generate snATACseq data using the 10X Chromium platform according to the manufacturer’s protocols (Demonstrated Protocol Nuclei Isolation for ATAC Sequencing Document CG000169; Chromium Single Cell ATAC_User Guide RevB Document CG000168). Briefly, >100,000 cells of interest were centrifuged, the supernatant was removed without disrupting the cell pellet, Lysis Buffer was added for 5 minutes on ice to generate isolated and permeabilized nuclei, followed by quenching by dilution with Wash Buffer. After centrifugation to pellet the washed nuclei, Diluted Nuclei Buffer was used to re-suspend nuclei at the desired nuclei concentration as determined using a Countess II FL Automated Cell Counter and combined with ATAC Buffer and ATAC Enzyme to form a Transposition Mix. Transposed nuclei were immediately combined with Barcoding Reagent, Reducing Agent B and Barcoding Enzyme and loaded onto a 10X Chromium Chip E for droplet generation, followed by library construction. The barcoded sequencing libraries were subjected to bead clean-up and checked for quality on an Agilent 4200 TapeStation, quantified by qPCR (KAPA Biosystems Library Quantification Kit for Illumina platforms), and pooled for sequencing on an Illumina NovaSeq 6000 S2 flow cell (paired-end libraries 2×50bp).

### Human islet isolation

Human islets were obtained through partnerships with the Integrated Islet Distribution Program (IIDP, http://iidp.coh.org/). Assessment of human islet function was performed by islet GSIS static incubation assay on the day after arrival, following the IIDP protocol. Primary human islets were cultured in Prodo media (PIM-S + supplements PIM-G + PIM-ABS) in 5% CO2 at 37oC for ∼24 hours prior to beginning studies. In preparation of single cell suspension for 10x platform, human islets were dispersed with StemPro Accutase (Thermo Fisher Scientific) 1ml/1000IEq for 10min at 37oC. Islet single cell suspension was washed three times in PBS-0.03% BSA and cell number determined using Countess II FL Automated Cell Counter (Life Tech). Nuclei isolation for single cell ATAC sequencing was performed following the 10x protocol (https://assets.ctfassets.net/an68im79xiti/5g035d2ngCW1aB9DFqPphO/71445a59fb282ea273a866c26cb5d319/CG000169_DemonstratedProtocol_NucleiIsolation_ATAC_Sequencing_RevD.pdf, based on the OMNI nucleiprep by Corces et al.^13^).

### Identifying snATAC-seq loci with >2 reads

Position sorted paired-end read alignments from snATAC-seq data are compared to detect all loci with >2 unique reads per nucleus. To avoid instances where reads overlap due to technical reasons, we removed all read pairs that are marked using the following parameters in the HTSJDK^14^ library: 1) *ReadPairedFlag* = True, 2) *ReadUnmappedFlag* = False, 3) *MateUnmappedFlag* = False, 4) *SecondaryOrSupplementary* = False, 5) *DuplicateReadFlag* = False, and *ReferenceIndex* != *MateReferenceIndex* (i.e., read pairs map to the same chromosome). To reduce overlaps due to alignment errors, reads are excluded based on i) mapping quality scores less than or equal to 30, and ii) insert sizes (i.e., the end to end distance between 5’ and 3’ read positions) greater than 900bp (∼6 nucleosomes) in length.

To identify instances of >2 reads overlapping at any specific locus, all intervals are identified for which an overlap was observed for at least two valid read pairs. Reads defining each interval are then compared to one another to identify all subintervals that exceed the specified overlap threshold (i.e., 2). To efficiently identify these subintervals, for each subset, interval breakpoints were defined at the start and end positions of each paired end read. For each interval breakpoint, an integer value of 1 was assigned to all breakpoints originating from start positions, and −1 to all breakpoints originating from an end position. Interval breakpoints are then visited in start position sorted order to generate a cumulative sum based on the assigned values at each breakpoint. The cumulative sum indicates the total number of overlaps between two interval breakpoints and efficiently identifies all sub-intervals with a number of overlaps greater than the specified threshold.

Once all subintervals satisfying the threshold are identified for a subset of reads, the algorithm repeats this process for the remaining paired end read subsets. Each step is performed using a linear time algorithm (i.e., O(n), n is the number of total reads), with an additional O(log(m)) (m equals the number of nuclei) overhead for each read to identify their respective nucleus origin, resulting in O(n*log(m)) runtime. The runtime can be reduced to an expected O(n) runtime by instead using an appropriate hash function for cell identifiers/barcodes. Note that this algorithm assumes that reads are sorted beforehand and is otherwise superseded by time it takes to sort reads by their chromosome and start positions (i.e., O(n*log(n)).

### Detecting significant multiplets from snATAC overlap counts

Loci with >2 reads were first filtered using simple repeats, segmental duplications, repeat masker and blacklist regions obtained from UCSC Genome Browser^15^ and ENCODE^16,17^. Next, filtered regions from all nuclei were merged if they overlapped by at least one base pair. Using this unified list of loci, a binary matrix was generated where rows in the matrix represent loci with >2 reads for at least one nucleus, and the columns represent the individual cells within the sample. Values within the matrix were assigned to 1 if the cell and genomic region combination observed >2 reads overlapping, and 0 otherwise. From this matrix, multiplets can be detected using column sums (i.e., the total number of >2 read overlap instances for each nucleus) while repetitive element sequences can be inferred using row sums (i.e., the total number of cells observing >2 reads at the same locus).

The events of observing >2 reads overlapping within the same region for multiple cells or across multiple regions within the same cell can be modeled using the Poisson distribution. Occurrences of these events are independent, counted within set intervals (i.e., counting regions across the entire genome within cells or counting cells within the same genomic regions), are either present or not within these intervals, and have a constant average rate of occurring, satisfying the assumptions of the Poisson distribution. We therefore detected significant multiplets and inferred repetitive sequences using the Poisson cumulative distribution function, using respective mean row and column sum counts as the expected values to calculate Poisson probabilities. In this process, we first use Poisson probabilities to infer repetitive sequences where a significant number of nuclei observe >2 reads at the same genomic region. All inferred repetitive sequence loci are removed from further analysis. Next, we calculate the Poisson probability of observing more loci with >2 reads than expected in a nucleus(i.e., multiplets) using column sums. Poisson probabilities for both inferring repetitive sequence and multiplet detection were corrected using the Benjamini Hochberg procedure to adjust for multiple hypothesis testing. Repetitive sequence inferences and multiplets were predicted by selecting regions or cells with adjusted Poisson probabilities less than 0.01.

### Multiplet annotation pipeline

Detected multiplets are annotated using clusters identified for snATAC-seq samples, merging them with respect to specific cell types present in the cell population. In our study, PBMC clusters were merged to represent CD4+T, CD8+T, Natural Killer (NK), myeloid and B cells and islet clusters were merged to represent alpha, beta, delta and ductal cells. Marker peaks for all cell type clusters with at least 150 cells were identified with the FindMarkers function in Seurat^18^, using the logistic regression setting. For the sake of unison, the top 100 marker peaks are then identified for each cell type cluster based on Bonferroni adjusted p-value of average log fold changes.

To account for data sparsity in snATAC-seq data, aggregate read profiles are calculated for each cell and marker peak. Aggregate read profiles are found by taking average read counts for each cell’s 15 nearest neighbors using the top 50 singular value decomposition (SVD) components. The cumulative distribution function in R (i.e., ecdf) is then used to find the abundance of reads for each cluster’s marker peaks. Distribution scores represent the percent of each cell type’s accessibility profiles present within the cell. In order to distinguish multiplet types (i.e., heterotypic or homotypic) singlet profiles were calculated for each cell type in the sample. For each cell type’s singlet cells, abundance scores at every marker peak were averaged to find the representive abundance score profile for that cell type. Multiplets that have a profile close to their abundant cell type’s singlet profile were classified as homotypic. Euclidean distance was used to measure the similarity between the profiles of multiplets and singlets. Mixture models were then fitted to the distances with the Mclust R package^19^ to group the closeness of the multiplets to their corresponding cell type’s singlet profile. Multiplets in the group with largest distance to the singlet profile are considered heterotypic. Multiplets are then annotated using the top 1 (for homotypic) or 2 (for heterotypic) abundance scores.

### snATAC-seq nuclei clustering

To cluster nuclei from snATAC-seq data, we employed an in-house implementation (https://github.com/UcarLab/snATACClusteringPipeline) of a two pass clustering method previously described^3^ with notable differences. First, we restrict the number of 2.5kb bins in the first pass clustering to the top 50k bins, up from 20k bins. For second pass clustering, we increase the number of peaks to include all peaks identified in pass 1 up to 200k.

### Integration of scRNA-seq and snATAC-seq data

Integrative clustering and analysis of single cell transcriptomes and single nucleus epigenomes was performed using the R package Seurat^18,20^. First, gene activity scores were derived from the resultant snATAC-seq peak count-matrix using the CreateGeneActivityMatrix function with default parameters. Next, single nuclei with < 5,000 total read counts were discarded from analyses. The resultant single nuclei and gene activity scores were log normalized and scaled. Using the processed scRNA-seq data (also analyzed with Seurat), we identified anchors between the snATAC-seq gene activity score matrix and scRNA-seq gene expression matrix following the methodology described by Butler et al. (2018)^18^. After identifying anchors between the datasets, cell-type labels from the scRNA-seq dataset were transferred to the snATAC-seq dataset and a prediction and confidence score was assigned for each cell.

### Simulating artificial multiplets to measure multiplet detection performances

To measure recall for detecting multiplets, artificial multiplets were simulated by combining accessibility profiles of nuclei within each sample population tested. For each sample, cells were randomly selected equal to 5% of the total cell population and paired together to introduce artificial multiplets equivalent to 2.5% of the total population. Introducing 2.5% artificial multiplets ensured that they were not the majority compared to real multiplets (5-11% of cells across all samples) present in the data. Cell pairs were randomly reselected until they formed heterotypic, homotypic, or 1:1 ratio of heterotypic and homotypic multiplets based on cell type annotations. Simulations measuring the number of valid read pairs per nucleus did not have restrictions based on cell type and were selected based on read depth when stratifying by number of valid read pairs (i.e., Fig. 3c-d, Extended Data Fig. 6b) or completely at random (i.e., Extended Data Fig. 7). Once cell pairs were identified, artificial multiplets were introduced by generating modified barcode mappings (for ATAC-DoubletDetector) or barcodes in fragment files (for ArchR^7^), which assigned artificial multiplet reads to the same cell identifier (i.e., the first nucleus in the pair). Artificial multiplets were simulated 10 or 100 runs depending on the analysis.

## CODE AVAILABILITY

ATAC-DoubletDetector is provided as a user-friendly computational framework with documentation and source code freely available at: https://github.com/UcarLab/ATAC-DoubletDetector.

---

**Extended Data Fig 1:**
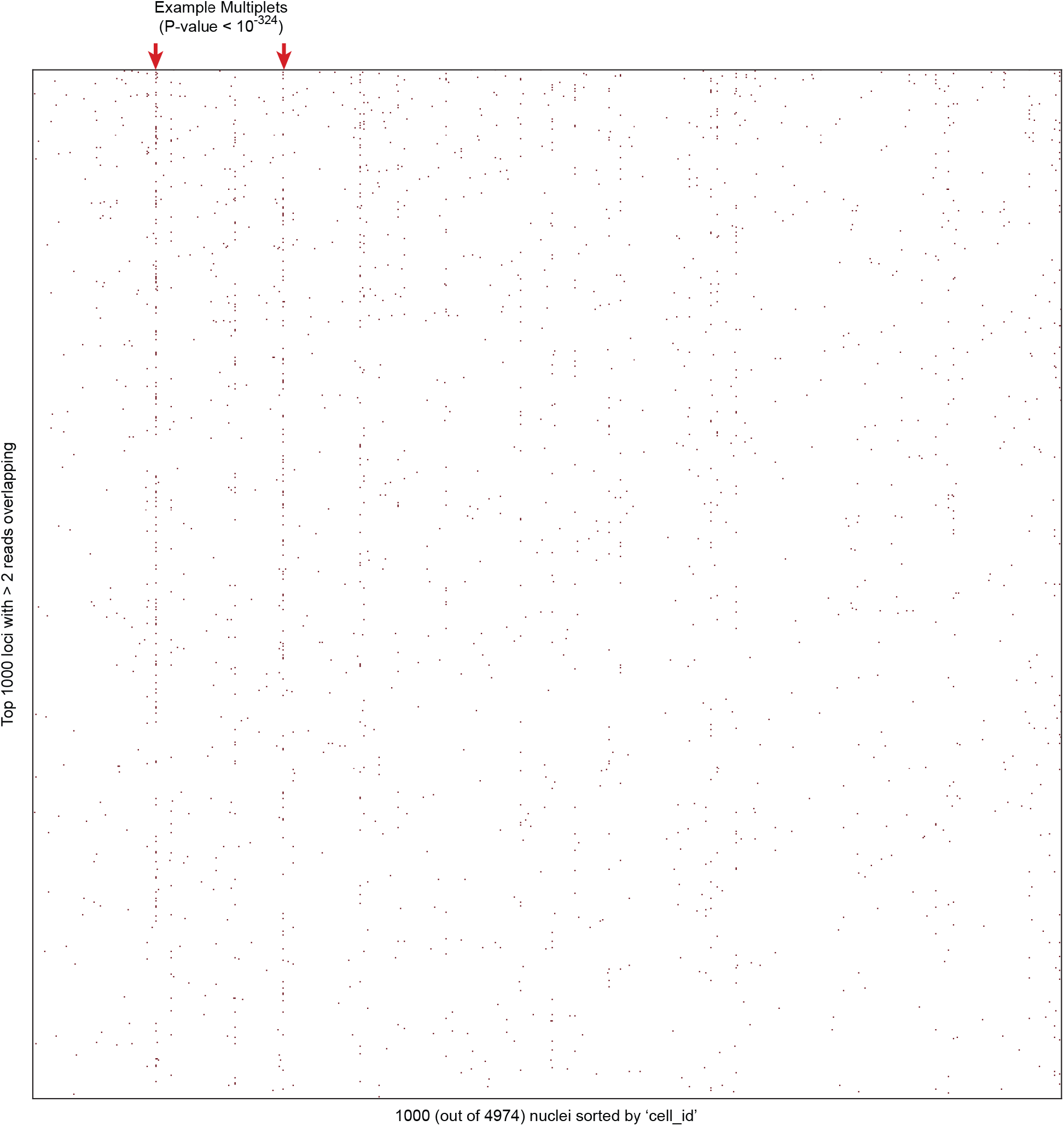
Multiplets observe many loci with >2 reads. The binary matrix of loci with >2 reads per cell reveals high confidence multiplet (marked by arrows) that harbor many loci with >2 reads. These multiplets can be clearly seen compared to the other cells in the subset.

**Extended Data Fig 2:**
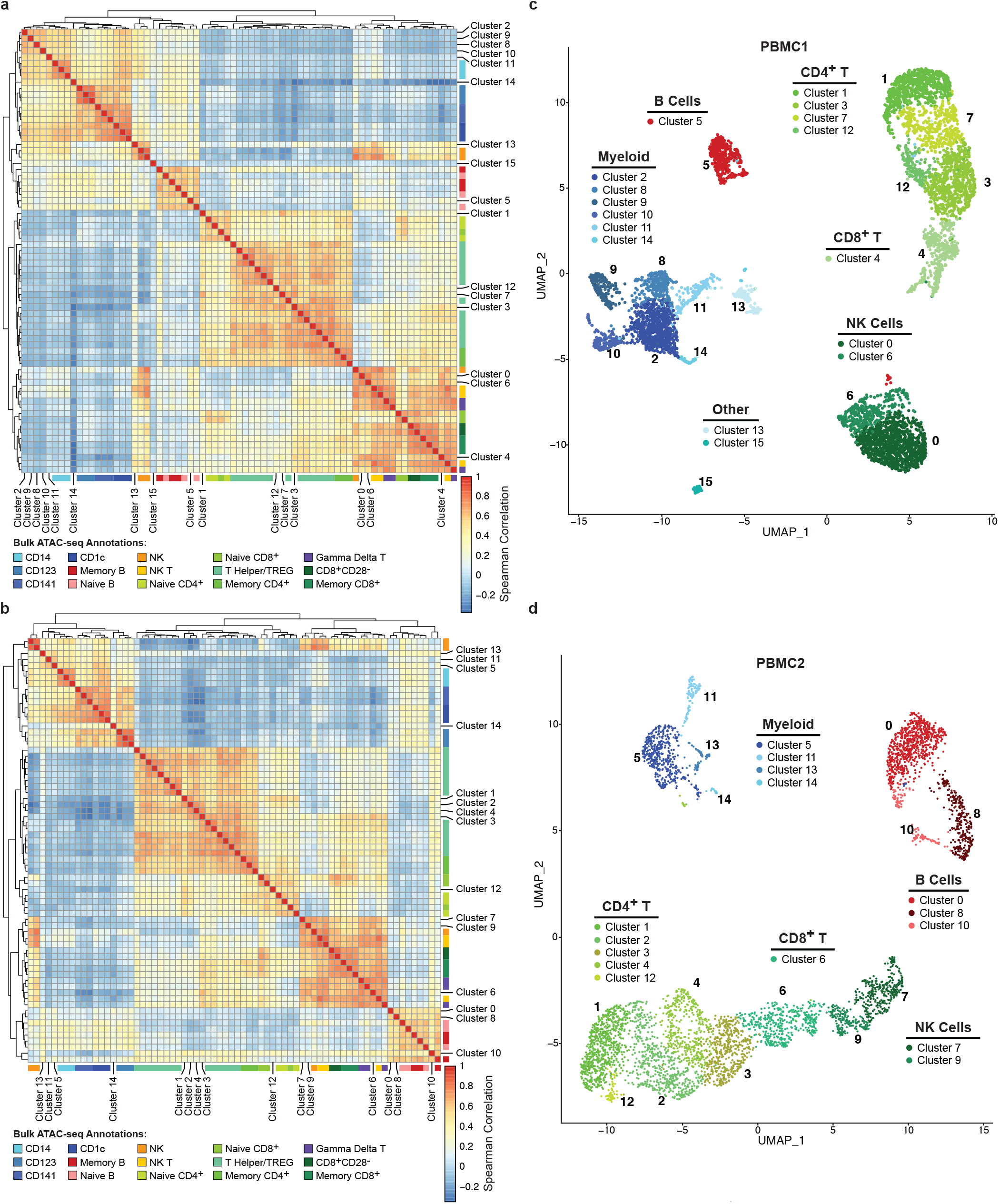
Pseudo-bulk snATAC-seq profile correlations with sorted bulk ATAC-seq revealed 5 major cell types. **a**,**b**, Spearman correlation heatmaps between pseudo-bulk (snATAC) and sorted bulk ATAC-seq accessibility profiles for PBMC1 (**a**) and PBMC2 (**b**). Pseudo-bulk profiles cluster with four major cell types: Myeloid, B, CD4^+^ T, CD8^+^ T and Natural Killer (NK). **c, d**, Annotated UMAP clusters for PBMC1 (**c**) and PBMC2 (**d**). Myeloid, B form distinct clusters for both samples. CD4^+^T, CD8^+^T and NK cell types share more accessible loci and tend to cluster more closely to one another.

**Extended Data Fig 3:**
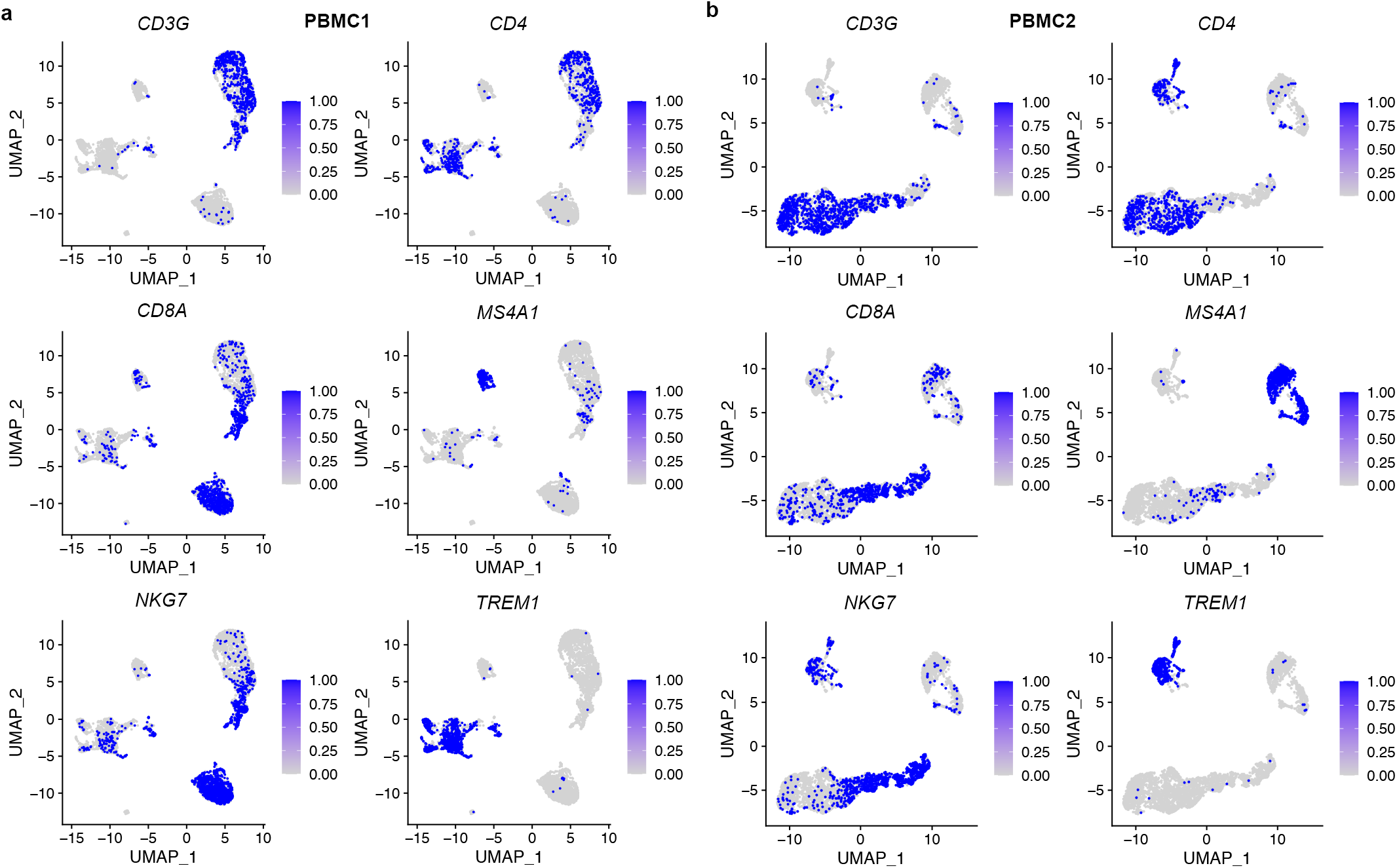
Annotated snATAC-seq clusters reflect accessibility at cell specific promoters. **a, b**, Annotated UMAPs for PBMC1 (**a**) and PBMC2 (**b**) at the promoters of *CD3G* (T-Cell Marker), *CD4* (CD4^+^ T cell marker), *CD8A* (CD8^+^ T cell marker), *MS4A1* (B cell marker), *NKG7* (NK cell marker), and *TREM1* (Myeloid cell marker). Accessibility was binarized to 0 or 1 based on the presence or absence of a read within these promoters. Using these markers, B and Myeloid cell types are clearly annotated with their respective markers. CD4^+^ T and CD8^+^ T cells can be observed by combining *CD3G* with *CD4* and *CD8A* markers respectively whereas NK cells are can be seen using *NKG7* and excluding nuclei with accessibility at *CD3G* promoter.

**Extended Data Fig 4:**
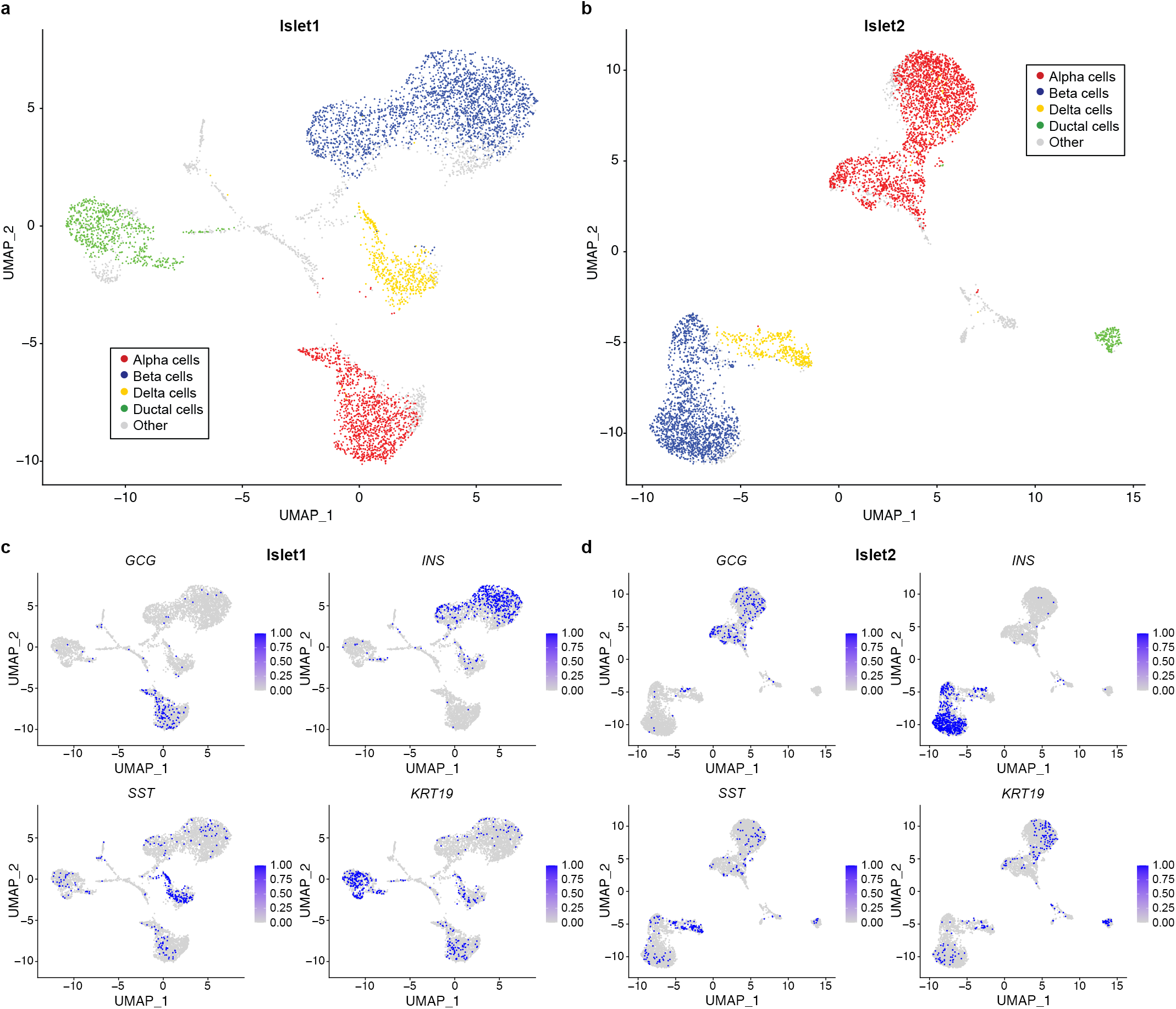
Islet snATAC-seq clusters correspond to scRNA-seq and cell marker annotations. **a, b**, UMAP clusters of snATAC-seq data for islet1 (**a**) and islet2 (**b**) annotated as alpha, beta delta or ductal cells *via* integration with annotated scRNA-seq data. Four distinct clusters are observed with these cell types. **c, d**. Cell specific clusters correspond to their respective marker peaks for both islet 1(**c**) and islet2 (**d**). Accessibility was binarized to 0 or 1 based on the presence or absence of a read within these promoters. Alpha, beta, delta and ductal cells are clearly identified with their respective marker genes: *GCG* (Alpha), *INS* (Beta), *SST* (Delta), and *KRT19* (Ductal).

**Extended Data Fig 5:**
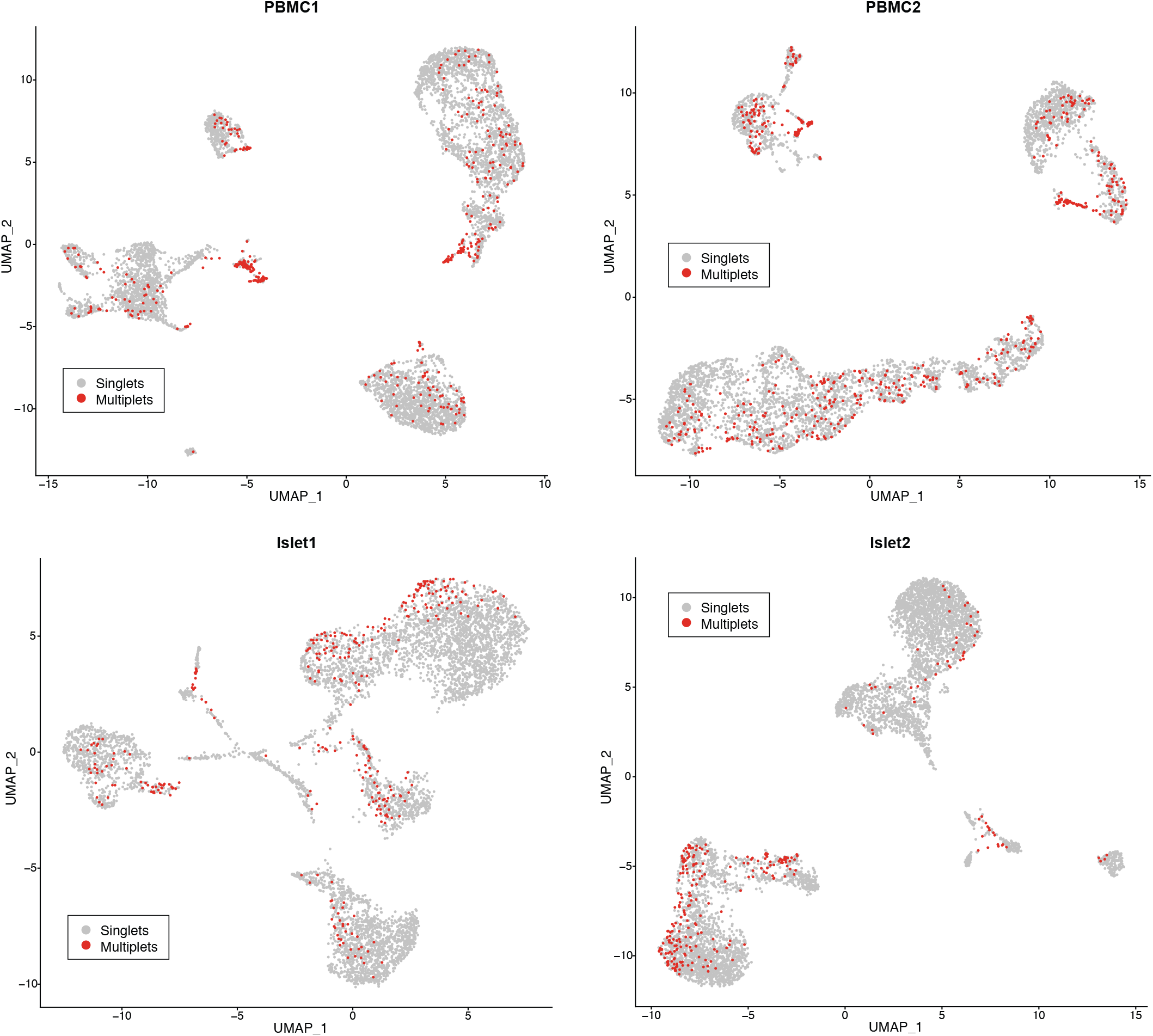
Multiplets are distributed throughout snATAC-seq clusters. Multiplet annotated UMAP clustering of PBMC1, PBMC2, islet1 and islet2 reveal that multiplets are distributed throughout all identified clusters and in some cases form their own multiplet clusters (i.e., center cluster in PBMC1). Multiplets between major cell type clusters are likely to be heterotypic whereas multiplets at the periphery of annotated clusters are likely to be homotypic.

**Extended Data Fig 6:**
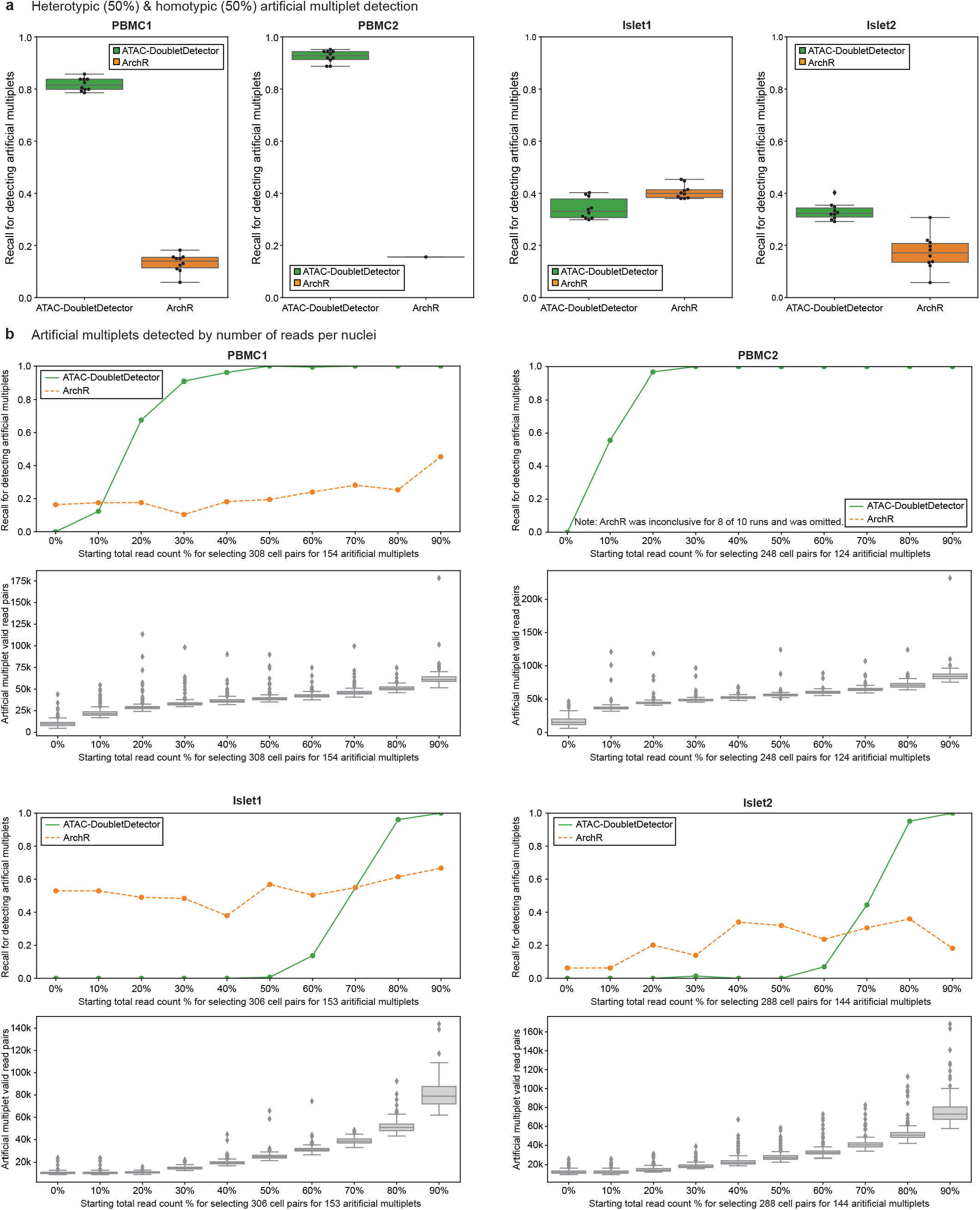
ATAC-DoubletDetector detects both homotypic and heterotypic multiplets at high read depth. **a**, Recall for detected both homotypic and heterotypic artificial multiplets at a 1:1 ratio. ATAC-DoubletDetector did not observe noticeable differences in performances due to its robustness for detecting both multiplet types. ArchR showed reduced performance compared to heterotypic multiplet only detection due to the inclusion of homotypic multiplets. **b**, Recall for multiplets stratified by read count distributions (top for each sample) and valid read pair distributions for each multiplet subset (bottom for each sample). ATAC-DoubletDetector performances increased when the number of valid read pairs exceeded ∼40k valid read pairs per nuclei, suggesting multiplets can be reliably detected when nuclei have >20k valid read pairs each. ArchR did not show significant differences in performance due to read depth.

**Extended Data Fig 7:**
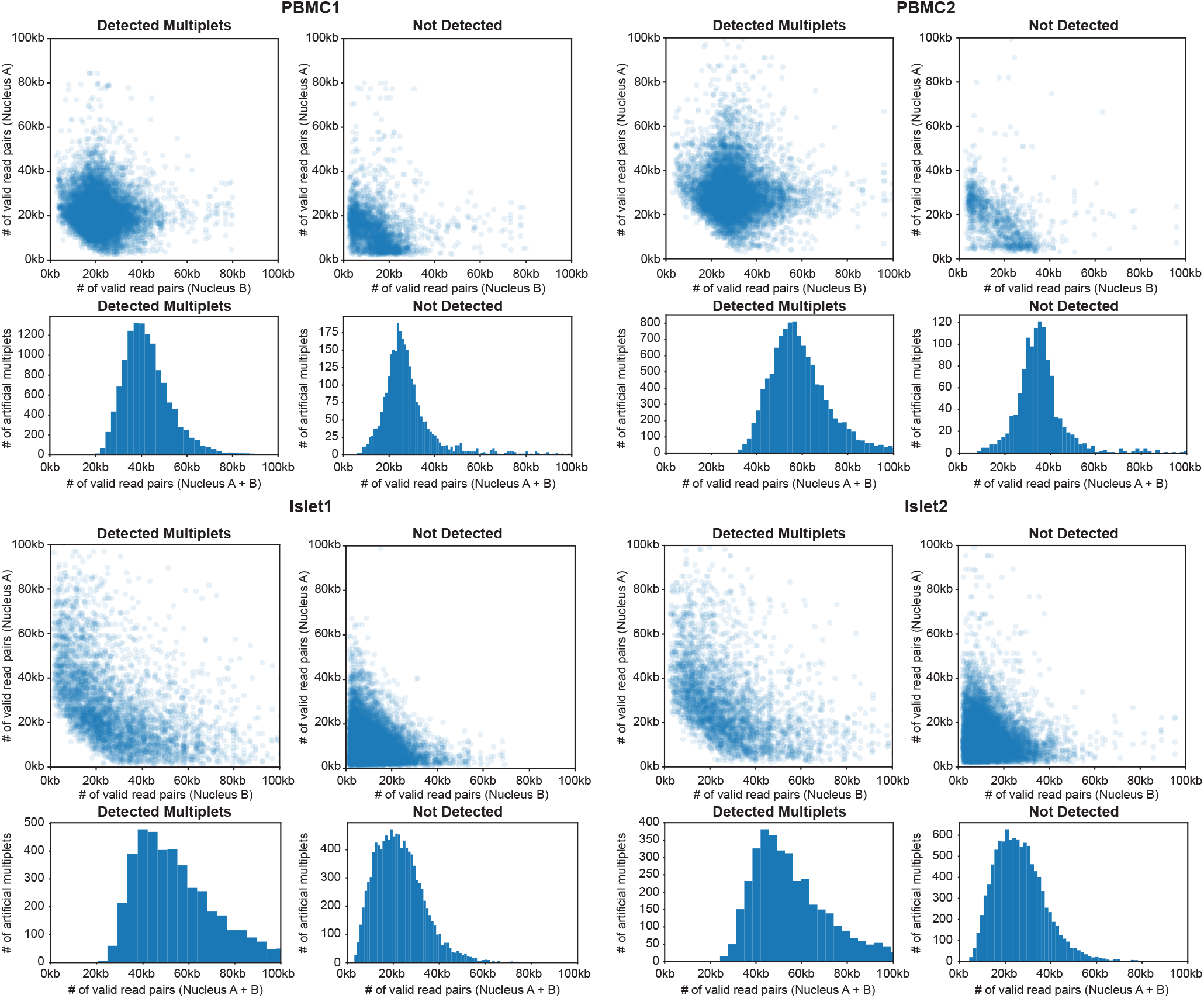
Artificial multiplets are detected when combined valid read pairs exceed 40k. For each sample, multiplets were detected (Top left for each sample) or not detect (Top right for each sample), depending on whether one or both nuclei exceeded 20k valid read pairs. Histogram of combined profiles revealed that the majority of detected multiplets (bottom left for each sample) had at least 20k valid read pairs while multiplets not detected were those with less than 40kb valid read pairs (bottom right for each sample). When nuclei are sequenced for 20k valid reads per nuclei, multiplets will harbor 40k valid read pairs and can be detected by ATAC-DoubletDetector.

**Extended Data Fig 8:**
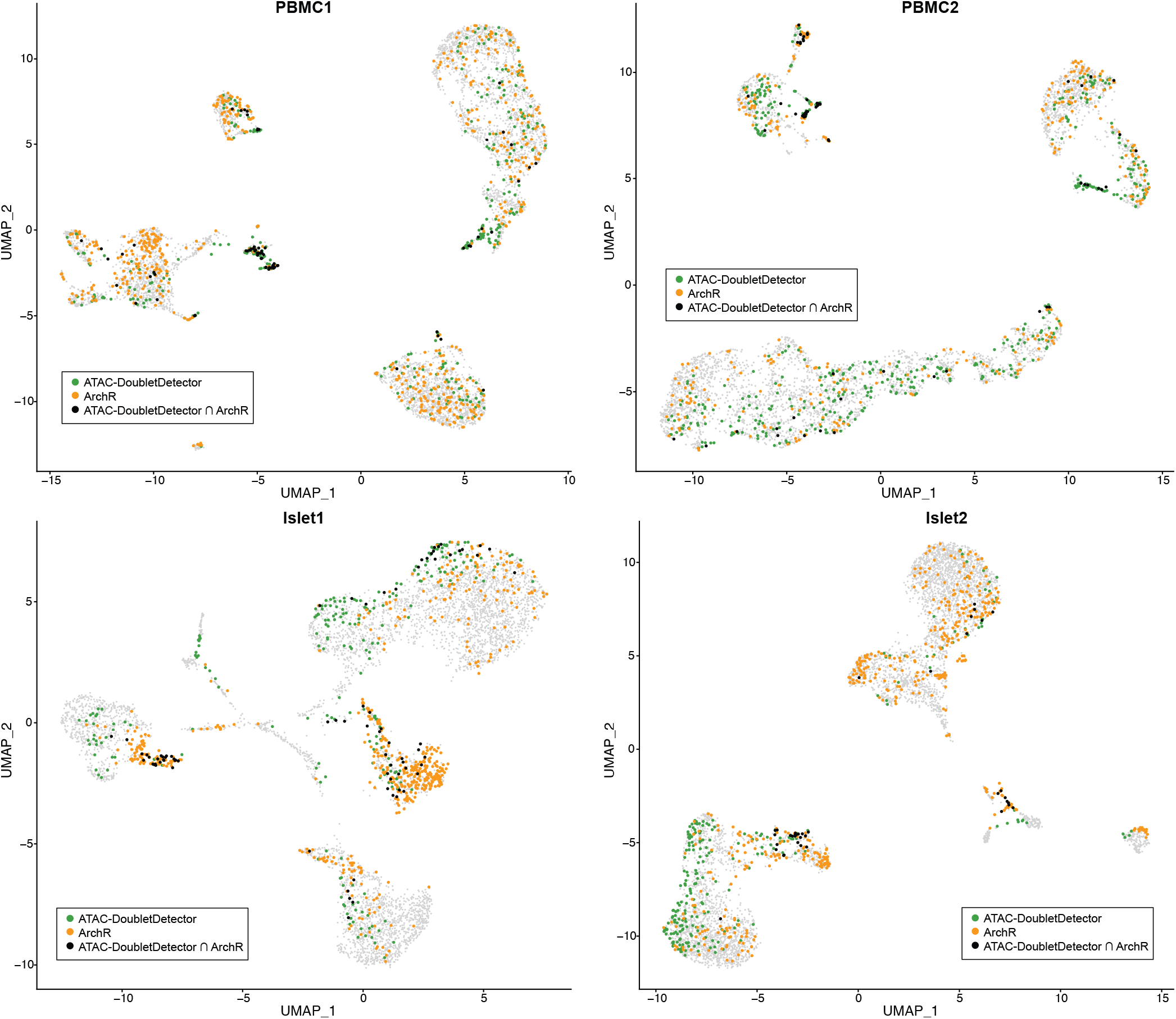
ATAC-DoubletDetector and ArchR identify different multiplet subsets. UMAP clusters annotating ATAC-DoubletDetector multiplets (green), ArchR multiplets (orange), or their intersection (black). Majority of multiplets detected by both ATAC-DoubletDetector and ArchR were between major cell type clusters (i.e., heterotypic multiplets).

**Extended Data Fig 9:**
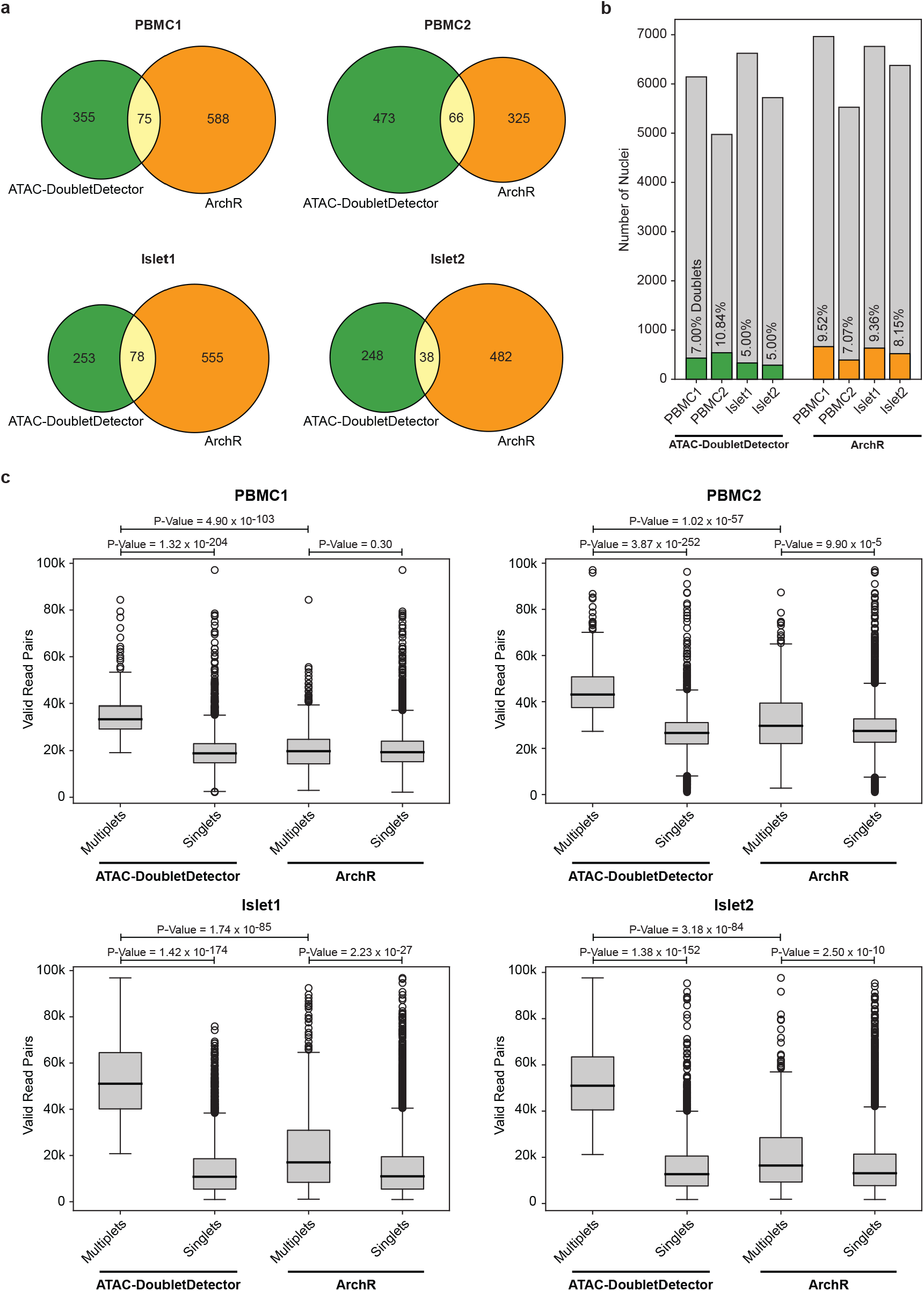
ATAC-DoubletDetector and ArchR multiplets comparisons reveal nature of their underlying algorithms. **a**, Venn diagrams and total number of multiplets detected by ATAC-DoubletDetector and ArchR. Only a small subset of multiplets is detected by both methods. **b**, Total number of nuclei and multiplets detected by each method. Differences in number of nuclei are due to differences in inputs (i.e., alignment (BAM) files for ATAC-DoubletDetector and fragment files (Cell Ranger output) for ArchR). Overall, ArchR detects more multiplets using default parameters than ATAC-DoubletDetector. **c**, Valid read pair distributions between multiplets and singlets detected by ATAC-DoubletDetector and ArchR. Differences in number of valid read pairs between multiplet and singlets were more significant for ATAC-DoubletDetector than ArchR while the number valid read pairs for ATAC-DoubletDetector were significantly greater than ArchR multiplet.

**Extended Data Fig 10:**
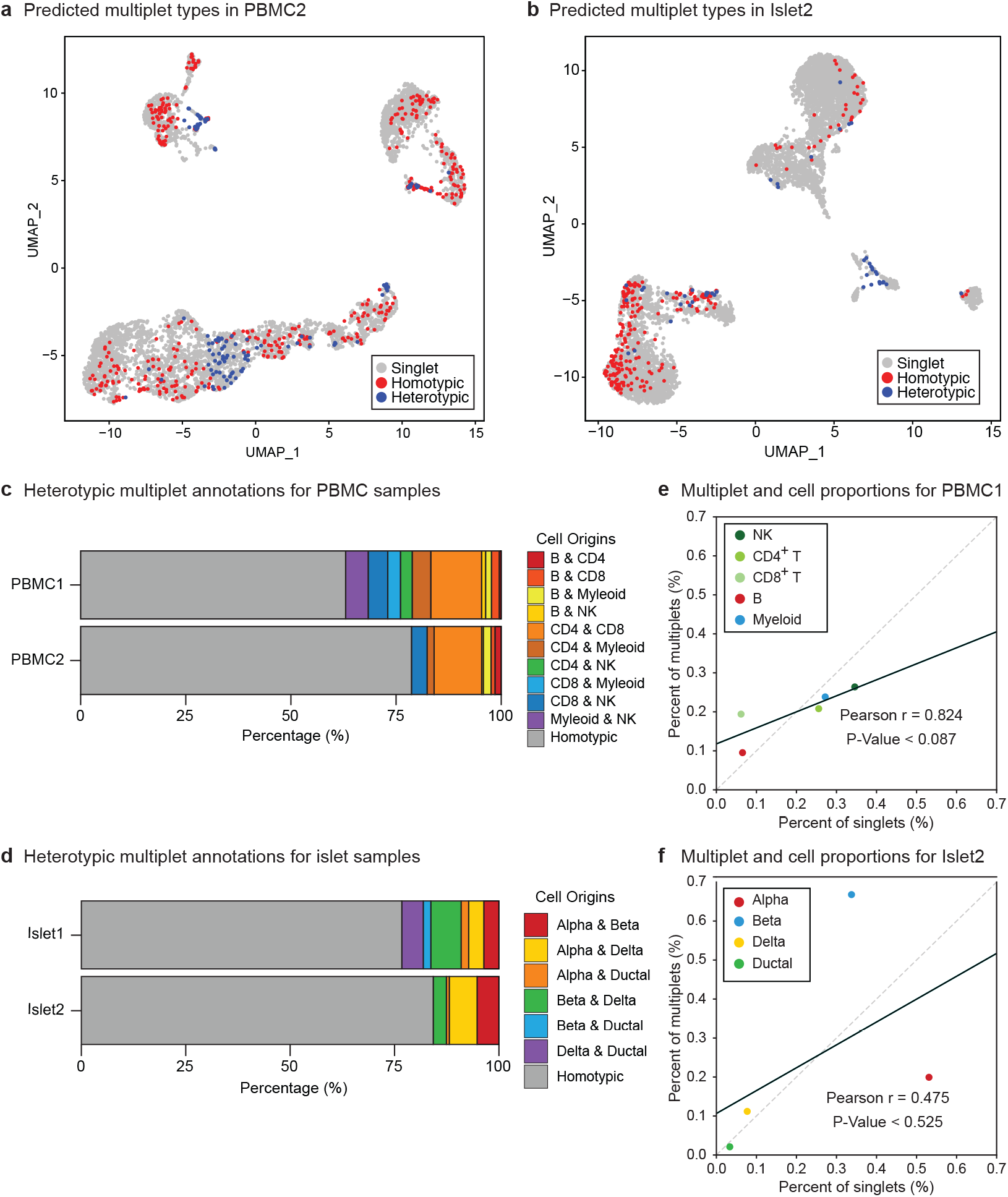
Multiplet annotations correspond to cell proportions. **a-b**, UMAP clustering for heterotypic and homotypic multiplet annotations in PBMC2 (**a**) and islet2 (**b**). Heterotypic multiplets are found between major cell type clusters. Homotypic multiplets are observed on the periphery of major cell type clusters. **c**-**d**, Heterotypic cell type annotations for PBMC (**d**) and islet (**e**) samples. Majority of multiplets are annotated as homotypic. **f**-**g**, Cell and multiplet proportions for PBMC1(**f**) and islet2(**g**). Multiplet cell type proportions are highly correlated with overall cell proportions. Islet2 observed more beta cell multiplets than other cell types/samples, reducing correlation and significance for islet2.

